# Exploring the limits of network topology estimation using diffusion-based tractography and tracer studies in the macaque cortex

**DOI:** 10.1101/356576

**Authors:** Kelly Shen, Alexandros Goulas, David Grayson, John Eusebio, Joseph S. Gati, Ravi S. Menon, Anthony R. McIntosh, Stefan Everling

## Abstract

Reconstructing the anatomical pathways of the brain to study the human connectome has become an important endeavour for understanding brain function and dynamics. Reconstruction of the cortico-cortical connectivity matrix *in vivo* often relies on noninvasive diffusion-weighted imaging (DWI) techniques but the extent to which they can accurately represent the topological characteristics of structural connectomes remains unknown. We explored this question by constructing connectomes using DWI data collected from macaque monkeys *in vivo* and with data from published invasive tracer studies. We found the strength of fiber tracts was well estimated from DWI and topological properties like degree and modularity were captured by tractography-based connectomes. Rich-club/core-periphery type architecture could also be detected but the classification of hubs using betweenness centrality, participation coefficient and core-periphery identification techniques was inaccurate. Our findings indicate that certain aspects of cortical topology can be faithfully represented in noninvasively-obtained connectomes while other network analytic measures warrant cautionary interpretations.

## Introduction

Network structure is thought to play a prominent role in supporting healthy brain function (Griffa et al., 2013; Fornito et al., 2015). Indeed, a large body of work has been devoted to the analysis of the brain’s structural topology in order to characterize and infer how functional networks emerge from large-scale structural connectivity, or the “connectome” (Park and Friston, 2013; Sporns, 2014; Zuo et al., 2016). In humans, the characterization of network structure relies mainly on noninvasive techniques such as tractography using diffusion-weighted magnetic resonance imaging (DWI). A number of influential observations about brain organization in both health and disease have been made based on DWI data (e.g., van den Heuvel et al., 2010; van den Heuvel and Sporns, 2011; Zalesky et al., 2011; Crossley et al., 2014; Perry et al., 2015; Baum et al., 2017). Recent validation studies in the macaque have demonstrated how a general correspondence exists between DWI-based estimates of structural connectivity, specifically “connection strength” (usually taken as some derivative of the number of streamlines between two regions), and those derived from the gold standard invasive technique of using tract tracers to map axonal projections. DWI-based tractography has been shown to correctly detect the presence of a large proportion of connections across the visual system (Azadbakht et al., 2015) and DWI-based estimates of connection strengths are correlated to those obtained from tracer studies (van den Heuvel et al., 2015; Donahue et al., 2016). However, even with extremely high-resolution DWI, probabilistic tractography suffers from a steep trade-off between sensitivity and specificity whereby obtaining a large proportion of true positive connections is accompanied by a large number of false positives and the optimal parameter settings for tractography (e.g., curvature thresholds) can vary widely depending on the location of the seed (Thomas et al., 2014; also see Maier-Hein et al., 2017). The ability of tractography to properly reconstruct the connectivity of the human brain and, in particular, the interpretation of detected streamlines (Jones et al., 2013), remains a matter of debate.

Existing validation studies in the macaque have only examined the accuracy of DWI-based tractography at the level of the individual connection. However, a major use of tractography has been to study the human brain at the level of large-scale whole-brain networks. The extent to which tractography can accurately capture the brain’s structural topology remains unknown. While some studies have shown that connectomes generated from tracer studies exhibit similar network organization principles as those reported using DWI data (e.g., Harriger et al., 2012; de Reus and van den Heuvel, 2013a), it is still unclear whether the topologies of networks obtained from the two different modalities actually coincide. Most previous studies have also been limited to tractography within a single hemisphere and usually using only a few *ex vivo* specimens, where DWI scans are of optimal quality and are not affected by artifacts such as motion or physiological noise. In this study, we used DWI data obtained from 10 macaque monkeys in combination with macaque connectivity described by published tracer studies to determine whether probabilistic tractography can accurately represent whole-brain structural topology *in vivo*. Given that tractography’s accuracy varies greatly as a function of its parameter settings (Dauguet et al., 2007; Jones et al., 2013; Thomas et al., 2014), we first systematically varied tractography parameters to determine the optimal settings for constructing whole-brain connectomes in the macaque. Using these optimized connectomes in conjunction with network analytic tools, we then determined the extent to which connectomes derived from DWI accurately captured the structural network characteristics of the macaque brain. We replicate previous findings that tractography can detect the presence and/or absence of connections above chance levels and can also provide reasonable estimates of connection strengths. In the macaque, more accurate connectomes were obtained by lowering the curvature threshold and discarding a small percentage of the weakest connections. However, owing to the high false positive rates in tractography-based connectomes, their ability to accurately capture critical aspects of structural topology was dependent on the robustness of the network analytic measure in question to misidentified connections.

## Results

Probabilistic tractography was performed using an FSL-based pipeline on diffusion-weighted magnetic resonance imaging data collected from 10 macaque monkeys at 7T. Two different parcellations, a single-hemisphere one (“Markov-Kennedy” (Markov et al., 2014)) and a whole-cortex parcellation (“RM-CoCo” (Kötter and Wanke, 2005; Bezgin et al., 2012)) were used and tractography parameters (angular threshold and distance correction) were systematically varied (see Materials and Methods). Tractography-derived connectivity matrices for various parameter combinations for an example subject are shown alongside the tracer-derived matrices in Figure 1. For the purposes of this paper, we use the term connection “strength” to refer to the number or proportion of axons running between two regions in the case of tract tracing data and the number or proportion of streamlines running between two regions for DWI-based tractography.

**Figure 1.**
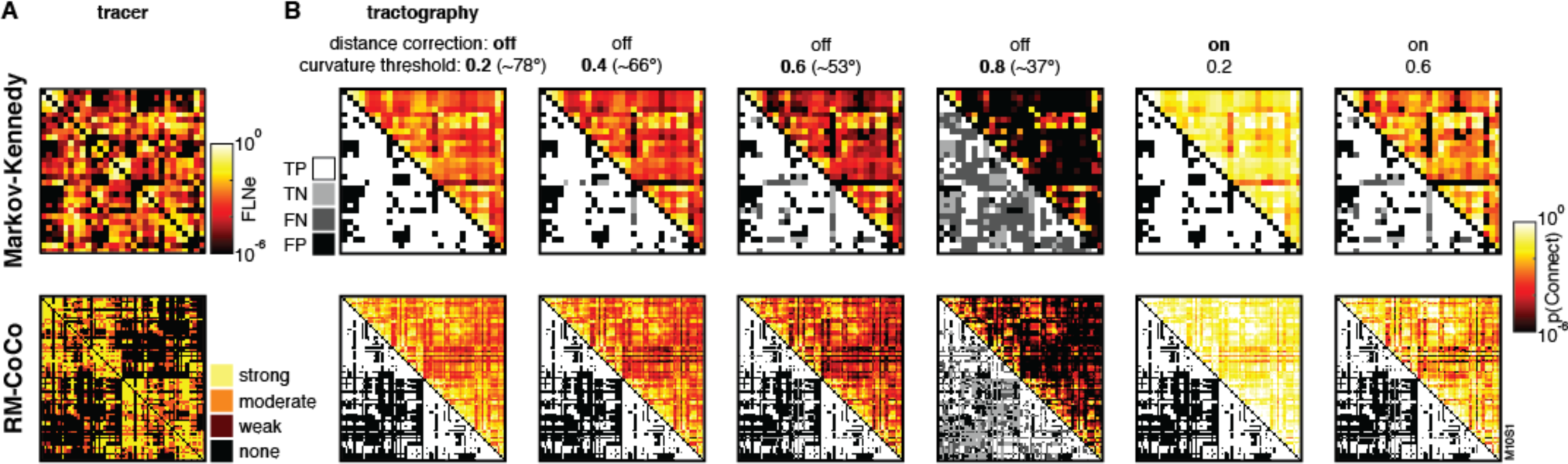
(A) Tracer-derived connectivity matrices from Markov et al (2014) (top) and CoCoMac (Stephan et al., 2001; Shen et al., 2012) (bottom). (B) Tractography-derived matrices (upper triangle) for an example subject for each parcellation (Markov-Kennedy, top; RM-CoCo, bottom) using various tractography parameters. Accuracy of each connection, as compared to tracer-derived matrices, depicted in lower triangles (TP: true positive; TN: true negative; FN: false negative; FP: false positive). For the RM-CoCo parcellation, left hemisphere ROIs are ordered together followed by right hemisphere ROIs, such that interhemispheric quadrants are the upper right and lower left of each matrix.

### Effects of varying tractography parameters on accuracy

On the assumption that the tracer-derived networks serve as a “ground truth” for the large-scale anatomical connectivity of the macaque brain, we computed a number of accuracy measures to determine the ability for diffusion-weighted tractography to reconstruct anatomical connectivity from data collected *in vivo.* These included the percentage of connections correctly represented, the area under the ROC curve (AUC), and corresponding measures of sensitivity, specificity and precision. We first consider intrahemispheric tractography using the Markov-Kennedy parcellation. With the “default” tractography parameter combination (curvature threshold: 0.2; distance correction: off), the percentage of connections correctly represented in the tractography-derived connectivity matrices was on average 79.21% (SD: 0.32) before any thresholding was performed. The mean AUC was 0.68 (SD: 0.02), which corresponded with a very high sensitivity (M: 0.99, SD: 0.01) but very low specificity (M: 0.01, SD: 0.01). These results are in line with previous macaque studies using *ex vivo* specimens that suggested that probabilistic tractography is accurate at correctly detecting connections (Azadbakht et al., 2015) but trades off specificity for sensitivity (Thomas et al., 2014). Precision was, on average, 0.79 (SD: 0.002) for the default parameter settings indicating a high positive predictive value in DWI tractrography (i.e., the great majority of positive results are true positives).

Curvature thresholds in tractography algorithms constrain the extent to which estimated streamlines can turn as they propogate. By default, FSL’s algorithm uses a threshold of 0.2, corresponding to ~78°. We systematically lowered this threshold (0.4, 0.6, 0.8 or ~66°, ~53°, ~37°) to examine its effect on the accuracy of tractography. There was an effect of curvature threshold on the percentage of correctly detected connections of the unthresholded matrices (i.e., where the x-axis = 0, Fig. 2A; repeated measures one-way ANOVA, F(3, 9)=525.85, p<0.001). Notably, post hoc comparisons indicated that % correct was not significantly different between matrices derived using curvature thresholds of 0.2 and 0.4 (M: 79.14%, SD: 0.12) but was significantly lower for thresholds of 0.6 (M: 76.21%, SD: 0.63) and 0.8 (M: 51.31, SD: 1.02) (Tukey-Kramer tests, p < 0.05). The effect of curvature threshold on the AUC of unthresholded matrices was limited to differences between the lowest threshold (0.8) and all other thresholds (repeated measures one-way ANOVA, F(3,9)=39.83, p<0.001; post hoc Tukey-Kramer tests) (Fig. 2B). Lowering the curvature threshold to 0.6 and below resulted in a significant drop in sensitivity (repeated measures one-way ANOVA, F(3,9)=676.26, p<0.001; post hoc Tukey-Kramer tests) with no differences for curvature thresholds of 0.2 and 0.4 (Fig. 2C, top). This was accompanied by a significant increase specificity across all curvature thresholds (Fig. 2C, bottom; repeated measures one-way ANOVA, F(3,9)=530.79, p<0.001; post hoc Tukey-Kramer tests). Precision also significantly increased when the curvature threshold was lowered to 0.6 and below (repeated measures one-way ANOVA, F(3,9)=81.26, p<0.001; post-hoc Tukey-Kramer tests), with no pairwise differences between 0.2 and 0.4 (Fig 2D). Intrahemispheric tractography using the RM-CoCo parcellation produced similar results (Fig. S1).

**Figure 2.**
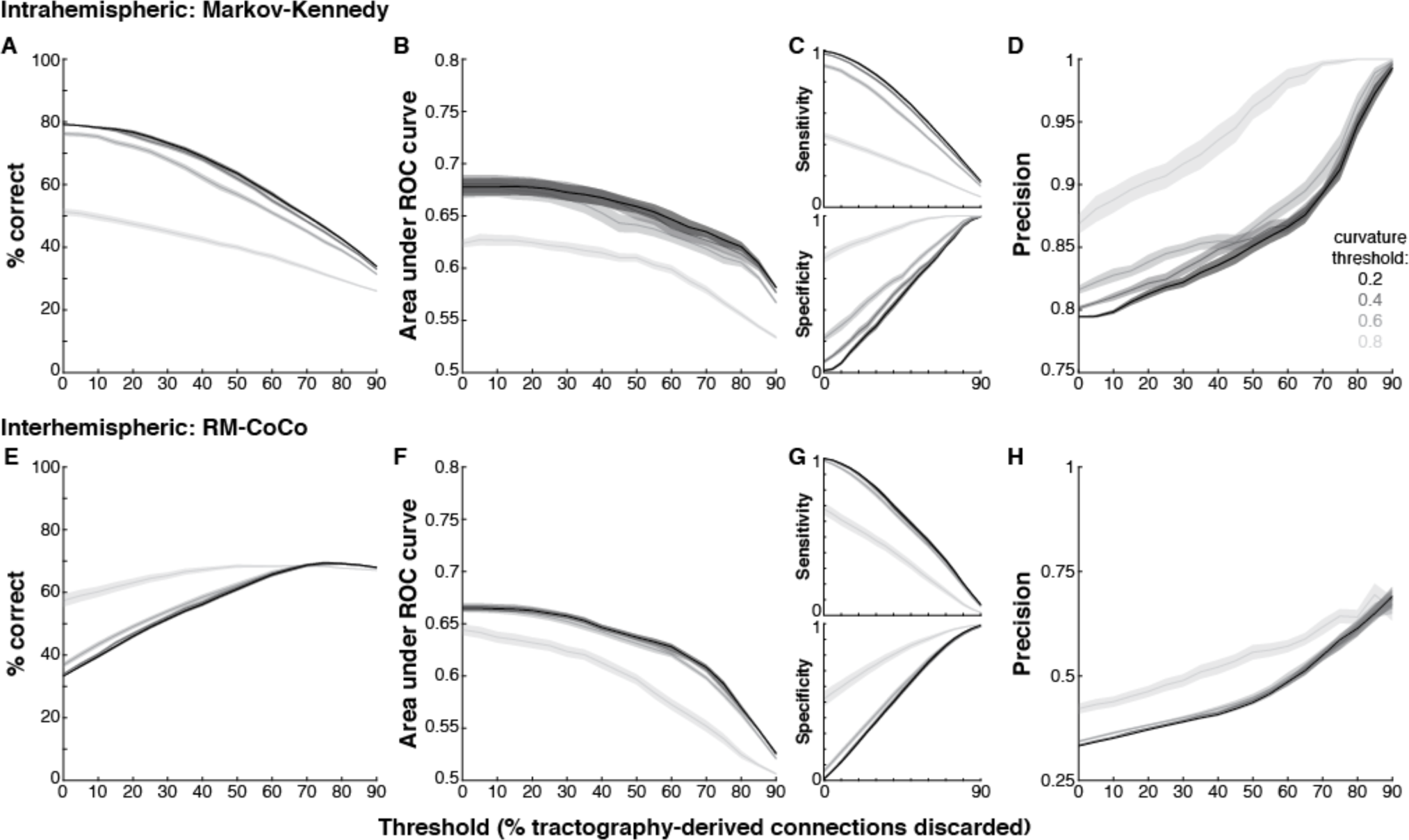
Accuracy of DWI tractography. (A-D) Accuracy measures for tractography using the Markov-Kennedy parcellation. (E-H) Accuracy measures for interhemispheric tractography using the RM-CoCo parcellation. Curves shown correspond to different tractography curvature thresholds as a function of thresholding the tractography-derived connectivity matrices (i.e., discarding connections having the lowest proportion of streamline counts).

We also explored the accuracy of interhemispheric tractography using the RM-CoCo parcellation. For the ‘default’ tractography parameter settings, interhemispheric tracking resulted in significantly lower % correct (paired t-test, t(9)=-1873.4, p<0.0001) and precision (t(9)=-5636.7, p<0.0001) while AUC was not different (t(9)= 1.74, p=0.12) (Fig. 2E-H vs. Fig. S1). However, as available tracer data for interhemispheric connections are limited, many of the “absent” interhemispheric connections in the tracer matrix are due to a lack of anatomical data. We therefore performed the same analysis on just the subset of interhemispheric connections for which the CoCoMac database indicates an explicitly present (n=479) or absent (n=125) connection. While accuracy of tractography was remarkably better for this subset of interhemispheric connections, it was still slightly lower than that of intrahemispheric tractography (Fig. S2). Of note, precision for the subset analysis was considerably higher (Fig. S2D) than that for all interhemispheric connections (Fig. 2H), indicating that the vast majority of detected connections in the explicit subset were true positives.

Just as in intrahemispheric tracking, there were no pairwise differences in % correct, AUC or precision values between curvature thresholds of 0.2 or 0.4 for interhemispheric tracking (repeated measures one-way ANOVAs, all p<0.001; post-hoc Tukey-Kramer tests) (Fig 2E-H). Together with the intrahemispheric tracking results, these findings suggest that using a high curvature threshold for macaque data does not result in a notable effect on the accuracy of DWI-based tractography and instead may lower the specificity when reconstructing anatomical connections across the macaque brain.

The ability of probabilistic tractography to reconstruct white matter fiber tracts is thought to be limited by the distance between ROIs. Factors such as noise, artifacts and actual fiber trajectory increase the uncertainty of tracking with increasing distance (Li et al., 2012). To test whether this was the case for our data, we binned connections by distance and found that % correct dropped as a function of distance for intrahemispheric tracking (Fig. S3A), consistent with previous findings for intrahemispheric tractography (Donahue et al., 2016). There was no consistent effect of distance on % correct for interhemispheric tracking (Fig. S3B). Employing distance correction did little, if anything, to change accuracy measures (data not shown), since our accuracy measures are computed using binarized data and distance correction as implemented in FSL is simply a reweighting scheme that biases the number of streamlines detected for long-distance tracts rather than whether streamlines are detectable.

Distance is also a determining factor in actual connectivity probabilities as observed in tracer-based networks (Markov et al., 2013; Beul et al., 2017), suggesting that the distance between ROIs could be used to estimate the existence of a connection between them. To test this, we used a simple geodesic distance-based model to generate connectivity matrices in the Markov-Kennedy parcellation. Remarkably, geodesic distance-based estimates of connectivity led to better correspondence with the tracer data (median AUC: 0.75) than DWI-based reconstructions (Fig. S4).

### Effects of discarding “weakest” connections on accuracy

For the purposes of connectome creation, the outputs of probabilistic tractography algorithms are often thresholded by discarding connections whose streamline counts do not meet a minimum requirement (Zalesky et al., 2016; Roberts et al., 2017). To determine whether such a thresholding technique improves the accuracy of probabilistic tractography, we systematically thresholded our tractography-derived connectivity matrices by discarding between 5 and 90% of the “weakest” connections (i.e., those with lowest weights) in increments of 5%. Accuracy, as measured by % connections correctly detected and AUC, dropped as a function of thresholding the intrahemispheric connectivity matrix (Fig. 2A-B and S1A-B), with a significant drop occurring once 20% or more of the weakest connections were discarded, depending on the accuracy metric, parcellation and curvature threshold used (Table S1). For interhemispheric tractography, only AUC dropped as a function of discarding the weakest weights, corresponding to a drop in sensitivity (Fig. 2F-G, but see Fig. S2), with a significant drop occurring once a 35% or more was reached (Table S2).

### Do DWI-based connectomes accurately depict network characteristics?

To determine whether network characteristics were accurately captured in DWI-based connectomes, we first constructed an average DWI-based connectome for each parcellation using the set of “optimal” tractography parameters and thresholds that maximized AUC for each animal (see Methods, Fig. S5 and Table S2). We also included for analysis a DWI-based structural network in the RM-CoCo parcellation derived from control animals recently described by Grayson and colleagues (2017). This independent average network was constructed using different imaging sequences, image preprocessing and probabilistic tractography procedures from those described in the present paper. For replication purposes and to examine the dependence of our results on our specific sample and DWI sequences, we present analyses on this additional DWI-based network in the supplementary materials (see Figures S6-S8).

The edge weights of the average DWI-based network were correlated with those from the tract-tracing one for the Markov-Kennedy parcellation (Fig 3A; Spearman rank correlation coefficient (r_s_) = 0.51, p<0.0001) and also for the RM-CoCo parcellation (Fig 3B; r_s_ = 0.45, p<0.0001; also see Fig. S6B). This correlation is in line with (Donahue et al., 2016) or better than (van den Heuvel et al., 2015) previous studies, and suggests that the strength of a white matter fiber tract is well captured by tractography-based estimates. As distance correction biases the number of streamlines detected for long tracks, and therefore biases the weights of our DWI-based connectomes, we additionally constructed distance-corrected versions of the average DWI-based network for comparison with the tracer networks. The correlation between the tractography-based and the tract-tracing-based edge weights was worse than the correlation obtained without distance correction for both the Markov-Kennedy (r_s_ = 0.46, p<0.0001; Fig. S7A) and RM parcellation (r_s_ = 0.40, p<0.0001; Fig. S7B). Finally, because connection strength varies as a function of distance in both tracer and tractography data (e.g., Donahue et al., 2016) we also computed the partial correlation between tracer and tractography weights while controlling for distance between ROIs. The correlation between the weights was reduced by half for the Markov-Kennedy parcellation (r_s_ = 0.21, p<0.0001) but only marginally for the RM-CoCo parcellation (r_s_ = 0.43, p<0.0001).

**Figure 3.**
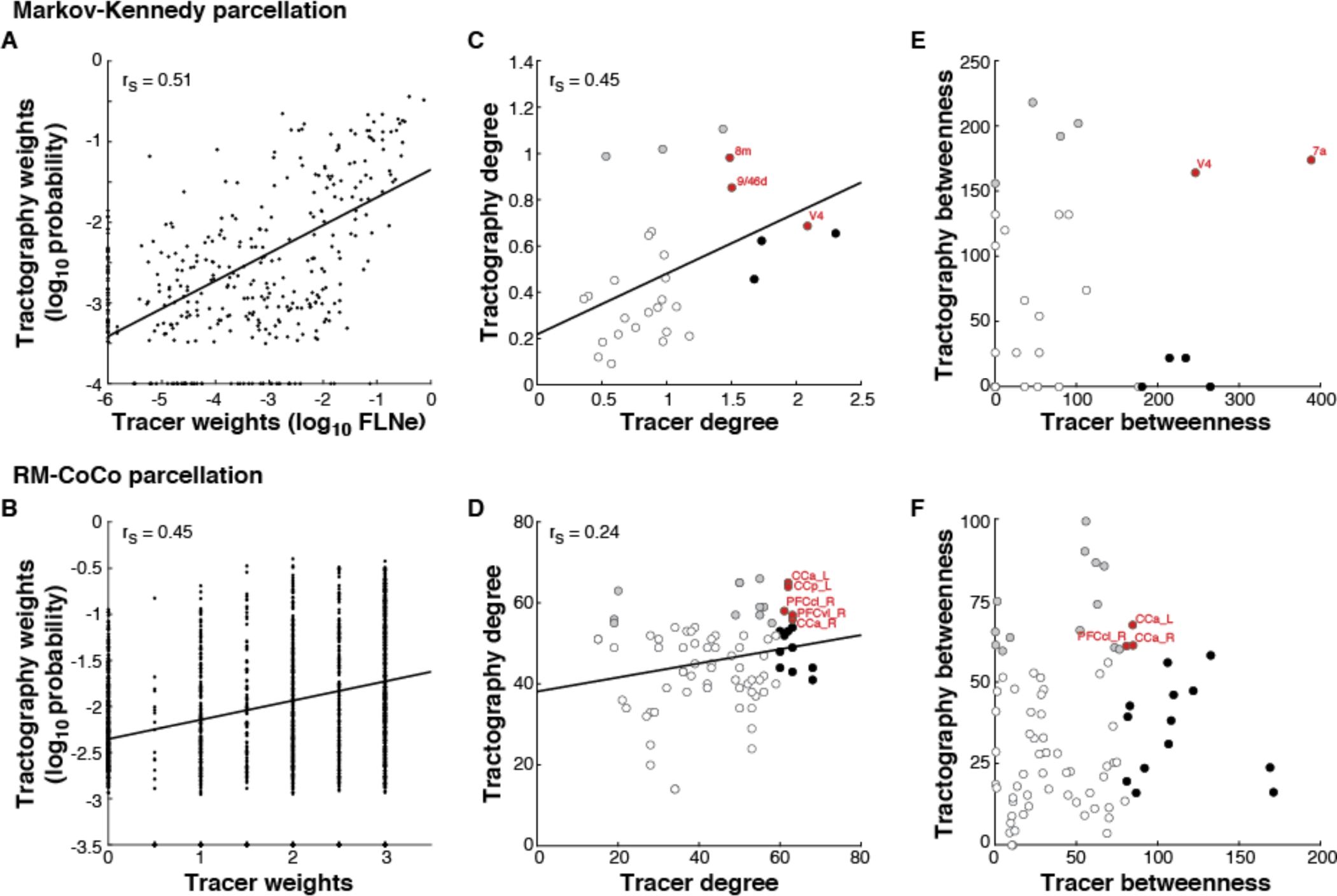
Correspondence of network topology metrics of ***in vivo*** tractography and tract-tracing connectomes. (A-B) Connectome weight estimates from DWI tractography are well correlated with those from tracer studies. (C-F) Centrality estimates from DWI-based networks are correlated with those from tracer studies for degree but not betweenness centrality estimates. Only some hubs, identified as those with centrality >80^th^ percentile, in the DWI-based networks correspond with hubs in tracer-based networks (red data points). These included cortical areas 7a, 8m, 9/46d, V4 in the Markov-Kennedy parcellation (C,E), and anterior and posterior cingulate cortex (CCa and CCp) and centrolateral and ventrolateral prefrontal cortex (PFCcl and PFCvl) in the RM-CoCo parcellation (D, F). Hubs in tracer-based networks not identified as hubs in DWI-based networks denoted in black; misidentified hubs in DWI-based networks that are not hubs in tracer-based networks denoted in grey. Correctly identified hubs denoted in red.

We next computed a number of graph metrics that capture different levels of description of topology for both tracer- and DWI-based networks to determine the extent to which tractography-derived networks can accurately estimate network topology.

### Centrality

Centrality measures are commonly used to provide estimates of the extent to which each node is embedded within a network, describing its potential contribution to network communication (van den Heuvel and Sporns, 2013a). Figure 3 shows how the nodal degree for DWI- and tract-tracing-based networks was positively correlated for the Markov-Kennedy intrahemispheric parcellation (Fig 3C; r_s_ = 0.45, p < 0.01) as well as the RM-CoCo whole brain parcellation (Fig 3D; r_s_ = 0.24, p = 0.01). Betweenness centrality, however, was not correlated for either parcellation (Markov-Kennedy: r_s_ = 0.14, p=0.23; RM-CoCo: r_s_ =0.14, p=0.11) (also see Fig. S6C). Network “hubs” are often singled out for investigation because of the special topological role they are thought to play in network communication and are identified as those nodes with high centrality. To determine whether hubs in the tractography-based networks coincide with those in the tracer-based networks, we identified hub nodes as those having centrality values greater than the 80^th^ percentile for each centrality measure. Although some overlap exists in the identified hubs from tractography- and tracer-based networks (Fig. 3C-F; red data points), a number of hubs in the tracer-based networks were not considered hubs in the tractography-based networks (Fig. 3C-F; black data points) and vice versa (grey data points; also see Fig. S6C-D). These findings suggest that tractography-based estimates of node centrality may not accurately reflect actual topologically central cortical regions.

### Network architecture

#### Modularity

One common way to describe brain network architecture has been to decompose brain networks into smaller communities or modules that are responsible for more specialized functions, and the connections between communities as serving the potential to integrate across these functions (Meunier et al., 2010; Sporns and Betzel, 2016). We examined whether the modular organization of tractography-based networks accurately reflected those obtained from tract tracing. Tractography-based networks showed a remarkably similar organization of subnetworks as compared to the tracer-based networks (Fig. 4A). For the Markov-Kennedy parcellation, the modular organization differed only in its assignment of three nodes (F2, F5, ProM) to the prefrontal module in the tractography-based network (Fig. 4A top right, dark blue) rather than a more fronto-parietal one in the tracer-based network (Fig. 4A top left, light green). However, even in the tracer-based network, these three nodes exhibit extensive and strong connectivity with the prefrontal module (Fig. 4A top left, grey edges). For the RM-CoCo parcellation, the decomposition of the tractography-based network resulted in a fourth module (Fig. 4A, bottom right, dark blue) and the assignment of some prefrontal areas (PFCdl, FEF, PMCm, PFCvl, PMCvl) to a more prefrontal module (light blue) rather than a more fronto-parietal one as in the tracer-based network (Fig. 4A, bottom left, light green). We determined the distance between the two sets of modules by computing the variation of information (VI) between them (Meilǎ, 2007). For both parcellations, the VI was significantly lower between the tractography-based partitions and the tracer-based ones as compared to the tracer-based null networks (Markov-Kennedy: 0.13 vs 0.70 ± 4.2×10^−15^; RM-CoCo: 0.27 vs 0.35 ± 3.0×10^−15^), suggesting that the tractography- and tracer-based partitions were more similar to each other than expected by chance.

**Figure 4.**
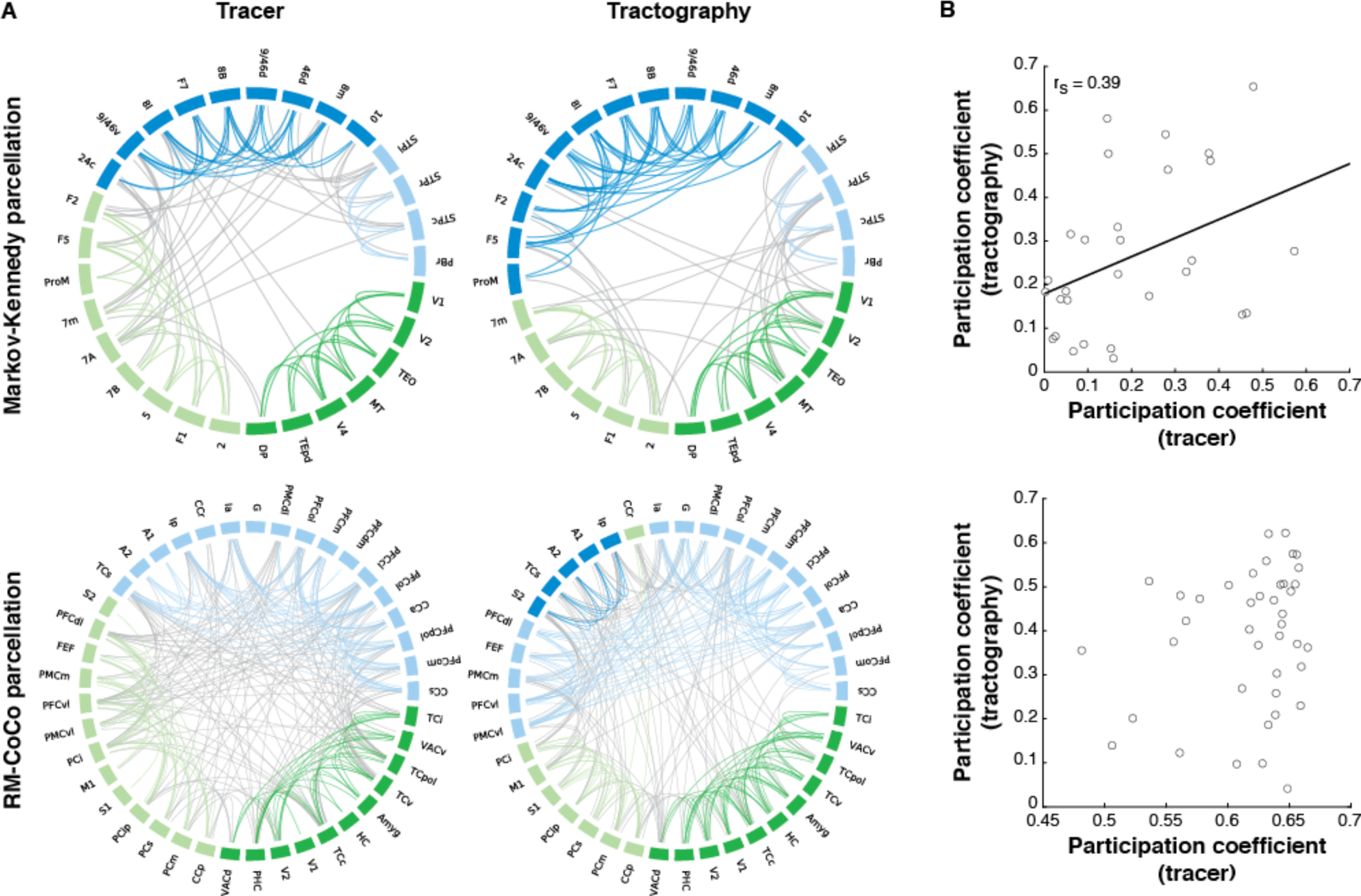
Modularity partitions of tractography-based networks are well matched with those from tracer-based networks. (A) Connectogram depictions of modular structure of each network type for the Markov-Kennedy (top) and the RM-CoCo (bottom) parcellations. Community assignments are denoted by node color. Within-module edges are denoted with the same color as the module while between-module edges are denoted in gray. Edges were thresholded for ease of visualization. For the Markov-Kennedy parcellation, the visualization threshold was set to 33% for both network types. For RM-Coco, the visualization threshold was set to keep connections in the strongest weight category (i.e., 3) in the tracer-based network and the tractography-based network was matched for the number of edges. (B) Correlation between tractography- and tracer-based network participation coefficients for the Markov-Kennedy (top) and RM-CoCo (bottom) parcellations. The community assignment of tracer-based networks were imposed on the tractography-based networks to determine participation coefficients.

We also assessed the accuracy of the tractography-based networks community assignments by first imposing the community structure of the tracer-based networks on the tractography-based ones and then computing the participation coefficients for each of the nodes in the tractography-based networks. The participation coefficient describes the extent to which each node is connected to nodes in other modules. If the “true” community assignments and, in particular, the between-module distribution of connections from the tracer-based networks were well estimated by the topology of the tractography-based networks, then the participation coefficients of the tractography-based network nodes computed in this manner should match those from the tracer-based networks. There was a moderate match for the Markov-Kennedy parcellation (Fig. 4B; r_s_ = 0.39, p=0.02, Spearman rank correlation) but not for the RM-CoCo parcellation (Fig. 4B; r_s_ = 0.03, p=0.42; also see Fig. S8). Together with the observed low VI between partition lists, these results suggest that nodal community assignments are well represented by DWI-based connectomes and the distribution of connections between and within modules can, to some degree, be estimated as well.

#### Rich Club Architecture

Brain networks have also been described as having a so-called “rich club” architecture, whereby a subset of high degree nodes exhibit dense connectivity with each other, often poised to mediate intermodular communication and forming a strong anatomical core (van den Heuvel and Sporns, 2013b). We examined whether a rich club architecture could be detected in the tracer-based networks, and the extent to which the tractography-based networks were able to replicate such findings. As our networks were weighted, we computed the normalized rich club coefficient by considering network weights in addition to topology when generating the null models (Alstott et al., 2014) for both of the tractography-based networks as well as the Markov-Kennedy tracer network. Similar to previous findings (Knoblauch et al., 2016), the Markov-Kennedy tracer network approached a rich club architecture at a degree of 24 (p = 0.08 for weighted networks, p = 0.06 for mixed networks) but rich club architecture was not consistently detected across a range of degrees and by and large not significant for any of the types of network considered (Fig. 6A, left). Although the DWI-based Markov-Kennedy network showed an increase in the normalized rich club coefficient at high degree levels, none were significantly greater than 1 following FDR correction (Fig. 5A, right). As we have previously reported (Shen et al., 2015), the RM-CoCo tracer network exhibits a rich club architecture at multiple degree levels (Fig. 5B, left). The DWI-based RM-CoCo network also exhibits a rich club architecture at multiple degree levels for all three types of models considered (Fig. 5B, right). A hypergeometric test of significant rich club levels detected in this network when a mixed model was considered showed significant overlap with the tracer network in 3 of 14 levels (at k=55, p<0.01; at k=54, p=0.01; and at k=53, p=0.03; also see Fig. S9). For level k=55, 10 rich club hubs that included regions of the prefrontal and cingulate cortex were identified in both the tracer- and DWI-based networks (Fig. 5C; red). Some additional regions of the temporal cortex along with dorso-medial prefrontal cortex were RC hubs in the DWI-based network but not the tracer-based one (Fig. 5C; grey), while a number of parietal and prefrontal RC hubs of the tracer-based network were notably missing from the DWI-based network (Fig. 5C; black).

**Figure 5.**
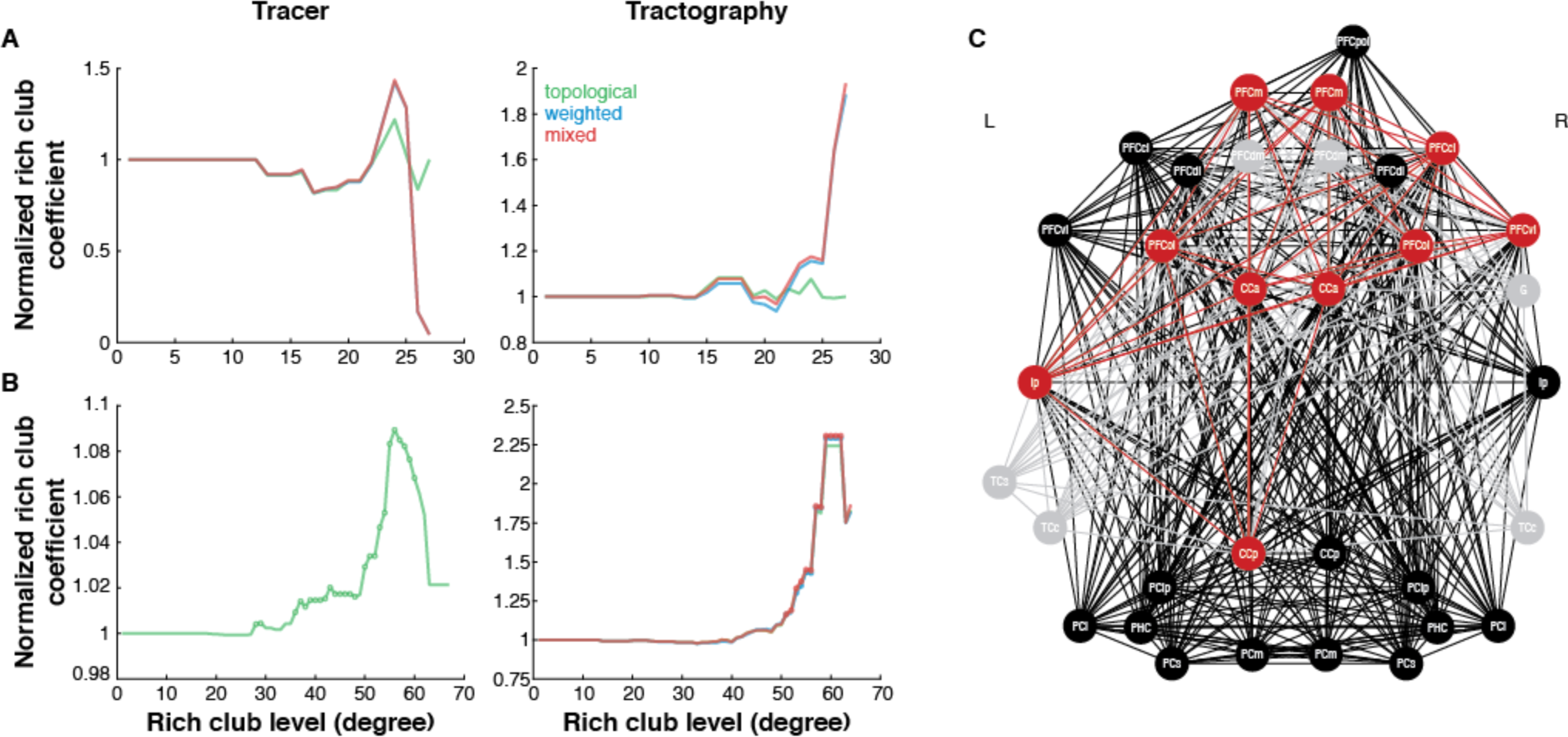
Rich club (RC) architecture in tracer- and DWI-based networks. Normalized rich club coefficient for tracer-(left) and DWI-based (right) networks for the Markov-Kennedy (A) and RM-CoCo (B) parcellations. RC levels (i.e., normalized rich club coefficients significantly >1) denoted by circles. (C) RCs at degree level 55 for RM-CoCo parcellation. Red nodes and edges depict those that are common to both tracer- and DWI-based networks. RC nodes and edges incorrectly detected by DWI are depicted in grey, and those in tracer-based networks but missed by DWI are depicted in black.

#### Core-Periphery Architecture

For denser networks, such as the Markov-Kennedy tracer one, a core-periphery architecture has been described (Ercsey-Ravasz et al., 2013). Here, we determined whether the core-periphery architecture previously reported in the Markov-Kennedy tracer network can be reconstructed by tractography-based connectomes. For the symmetrized tracer-based network, we detected a set of 18 nodes that contributed to the high-density core, with the 11 remaining nodes considered to be in the periphery (Fig. 6A). For the tractography-based network, a core-periphery architecture was also detected, with the core consisting of 21 nodes, of which 16 were correctly identified (Fig. 6B). A hypergeometric test showed significant overlap in the core memberships of the tracer and tractography networks (p<0.001).

**Figure 6.**
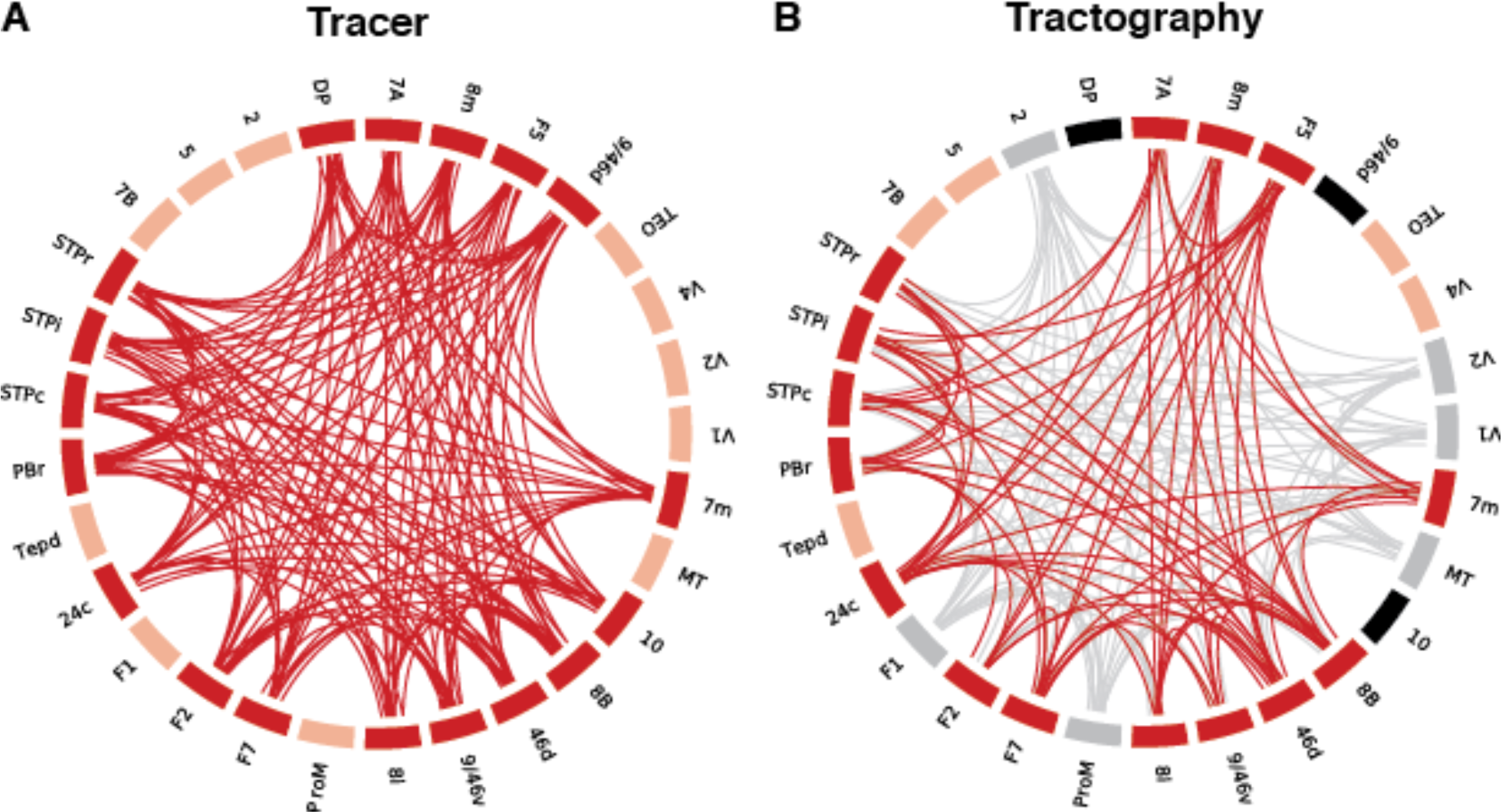
Core-Periphery architecture in tracer- and DWI-based networks. (A) Core-periphery architecture detected in Markov-Kennedy tract tracing network. Core nodes and edges depicted in red, peripheral nodes depicted in coral. (B) Coreperiphery architecture detected in the tractography-based network. Correctly identified core nodes and edges in red, peripheral nodes in coral. Incorrectly identified core nodes and edges depicted in grey, while core nodes that were missed depicted in black.

## Discussion

We have tested the performance of *in vivo* diffusion and tractography-based connectomes by comparing them to the gold standard connectomes from tracer data in macaque monkeys. We found that the reconstruction of individual connections to be moderately accurate, with a steep tradeoff between sensitivity and specificity that replicates previous *ex vivo* reports. We also found the proportion of streamlines detected between any two given regions can serve as a robust estimate of the number of axons that run between them, and that this relationship was dependent on the distance between regions. Importantly, we performed a series of validation studies on network topology metrics and demonstrated how the assignment of nodes into communities in **t**ractography-based connectomes is fairly accurate and that a high-density rich club or core-periphery organization can be detected, just as they can be in the corresponding tracer-based networks. However, the proper identification of hubs within modules, and membership in rich club or core-periphery type architectures was less accurate, likely owing to the great number of **f**alse positive connections generated by tractography. As network analysis has quickly become a popular approach for analyzing cortical connectomes, leading to the influential and expanding **f**ields of connectomics and network neuroscience (Sporns, 2013; Bassett and Sporns, 2017), our **f**indings are instrumental for the interpretation of network topology results based on *in vivo* measurements of structural connectivity.

### Reconstruction of interareal cortico-cortical connections *in vivo*

Mapping the cortical connectome and uncovering its topological layout is a major ongoing research endeavour, involving many large-scale efforts like the Human Connectome Project (Van Essen et al., 2013). Structural connectomes *in vivo* can only be constructed with diffusion weighted **i**maging and tractography at present. The majority of such approaches aim to reconstruct the large-scale cortico-cortical connectivity matrix and subsequently analyze it using network metrics (Bassett et al., 2008; Hagmann et al., 2008; Gong et al., 2009). Here, we found the reconstruction of connections between the cortical areas to be above chance, but not highly accurate. Our obtained quantitative measures, such as AUC, are comparable to recent reports investigating intrahemispheric connections (Thomas et al., 2014; Azadbakht et al., 2015; van den Heuvel et al., 2015; Donahue et al., 2016). Our results provide additional quantitative evidence on the feasibility to correctly uncover the correct pairs of interconnected areas with diffusion imaging and tractography, by using two different benchmark datasets obtained by tract tracing. Despite large differences in how these two benchmark datasets were derived, how their connectivity was expressed and how they have differing network topologies, intrahemispheric tractography results for these two parcellations were remarkably similar. Importantly, we also examined interhemispheric tractography and show that it exhibits worse reconstruction quality than intrahemispheric tractography. This is consistent with our observation that tractography accuracy decreases with increasing distance between regions (also see Li et al., 2012; Donahue et al., 2016), which is only compounded when tracking across hemispheres. However, when we limited our analysis to only those interhemispheric connections that were explicitly defined in CoCoMac, the accuracy of DWI tractography approached that of intrahemispheric tracking. Precision, in particular, was considerably higher for this subset analysis, suggesting that the false positive rate decreased dramatically when we only considered explicitly defined connections. This suggests that, to some degree, missing information in CoCoMac about interhemispheric connections may be lowering our estimates of DWI tractography accuracy. Moreover, although considered the “gold standard”, tracer data themselves are not perfect and can be affected by variability across individual injections, the uptake of tracers by passing axonal fibers, as well as the distance travelled by particular tracers (for discussion, see Köbbert et al., 2000; Lanciego and Wouterlood, 2011; Markov et al., 2014).

Earlier validation studies have highlighted the inability of tractography to resolve long-range connections. Both probabilistic (Li et al., 2012) and deterministic (Dauguet et al., 2007; Zalesky and Fornito, 2009) tractography algorithms suffer from false negatives associated with long-range fibers due to the increasing uncertainty of tractography with distance (Jbabdi et al., 2015). Simply biasing connection weights towards long-distance connections using the distance correction option did little to resolve this problem as accuracy measures relied on the binary classification of the existence or non-existence of connections (but see Azadbakht et al., 2015). Instead, implementing the distance correction option affected the ability of tractography to estimate the strength of connections, decreasing it substantially when distance was estimated with the more realistic measure of geodesic distance rather than Euclidean distance. Connectomes constructed using deterministic tractography are generally more sparse than those constructed using probabilistic tractography (Zalesky et al., 2016), and can suffer from a large number of false negative connections (Gong et al., 2009; Bastiani et al., 2012). Recent reports of probabilistic tractography, including the results presented here, have additionally indicated that while the majority of ‘true connections’ are successfully reconstructed, they instead come at the price of a large number of false positive connections (e.g., Thomas et al., 2014). The choice between deterministic and probabilistic tractography then, can be considered as a choice between constructing low-sensitivity/high-specificity connectomes versus high-sensitivity/low-specificity ones. Relevant for the probabilistic tractography results presented here, excessive false positive connections have been reported as a major drawback of diffusion imaging and tractography in a validation study with simulated human brain data (Maier-Hein et al., 2017). These findings, along with the observation that false positives have a much larger impact on estimates of network topology as compared to false negatives (Bastiani et al., 2012; Zalesky et al., 2016), should be explicitly taken into account as important limitations when interpreting results from diffusion imaging tractography. Our results indicate that thresholding the weakest weights in the tractography-based networks on the order of 20-30% did not affect the percentage of correctly detected connections. Moreover, thresholding on the order of 55-85% did not affect AUC as it decreased sensitivity while dramatically increasing specificity. This is consistent with previous estimates for optimizing the tradeoff between sensitivity and specificity (de Reus and van den Heuvel, 2013b; Donahue et al., 2016). Choosing to threshold by discarding the weakest weights, however, may result in also discarding weak true positives. Weak connections are known to be important in determining the brain’s functional organization (Gallos et al., 2012; Goulas et al., 2015 d may be better represented in networks that have been constructed using methods that take the frequency of edge reconstruction across subjects into account (Roberts et al., 2017). We also found that lower curvature thresholds, at least for macaque data, result in fewer false positives and greater specificity without greatly affecting other accuracy measures (also see Dauguet et al., 2007; Azadbakht et al., 2015). Whether this is a result of less cortical folding and, therefore, less convoluted white matter pathways in the macaque brain (Herculano-Houzel et al., 2010; Zilles et al., 2013) or whether it constitutes an indicative guide for human tractography remains to be seen.

Using the streamline count as a proxy of fiber density has been previously criticized because it is susceptible to differences in tract lengths, curvature and branching (Jones, 2010; Jones et al., 2013). However, we showed how the probability values obtained with tractography were significantly correlated with an explained variance in line with Donahue et al. (2016), and nearly twice that of van den Heuvel et al. (2015). Since it is clear from tracer studies that physical distance plays a large role in the existence and strength of connections (Markov et al., 2013; Beul et al., 2017), a cautionary note is needed when interpreting such validation results. We have shown that a model based on physical distance alone was able to achieve comparable and even higher AUC than the diffusion and tractography-based reconstructions. Moreover, the correlation between the strength of connections and the tractography probabilities was diminished when physical distance was taken into account. Physical distance-based models (e.g., Ercsey-Ravasz et al., 2013; Bezgin et al., 2017) may therefore offer a more stringent baseline than using tractography alone while advancements in both imaging and tractography methods are still needed for the accurate reconstruction of cortical connectomes *in vivo.*

### Investigating network topology of the cortex *in vivo*

The success of some network metrics but not others in the tractography-based connectomes was dependent on their resilience to rewirings. We found the partitioning of the cortico-cortical network into modules to be highly similar between the invasive and non-invasive connectivity datasets. These results bestow some confidence in module partitioning results obtained *in vivo.* This is in line with recent work that showed how false negatives, and even false positives, in connectomes affect modularity partitioning minimally (Zalesky et al., 2016). Partitioning brain networks into modules can result in variable communities across iterations (Sporns and Betzel, 2016). To minimize the effects of unstable partitionings, we chose to use the most consistent community structure detected from multiple iterations of partitioning. Additional work is still needed to fully assess how the accuracy of partitioning affects comparisons of modularity between different networks.

The participation coefficient, a higher-order network metric commonly used to identify intra-modular “provincial” and inter-modular “connector” hubs based on their cross-modular edges, was not consistent across the invasive and non-invasive datasets. Our results suggested that for the Markov-Kennedy parcellation, inaccuracies mostly arose from the reconstruction of inter- and intra-modular connections as the participation coefficients were correlated even when we controlled for differences in community structure by keeping the partitioning scheme fixed when computing participation coefficients. For the RM-CoCo parcellation, however, there was an additional cost from the slightly inaccurate classification of nodes into their respective communities. Indeed, two networks can have extremely similar modularity partitions but their underlying connections could be statistically independent.

The susceptibility of centrality measures to rewirings resulted in discrepancies between the results obtained from the invasive and non-invasive measurements. While for degree centrality a significant positive correlation was observed, no significant correlations were observed for betweenness centrality in our dataset and only moderate correlations in a second dataset (see Supplemental Material). The top most connected nodes or “hubs” between the two modalities also did not fully overlap when using either centrality metric. Our results indicate that more confidence can be assigned to degree as compared to betweenness centrality when non-invasive measurements are used. Along similar lines, global descriptions of structural organization like rich-club or core-periphery architectures, if they existed in the tract tracing networks, could be obtained from the tractography-based networks. However, the particular identification of nodes as hubs within these architectures was less accurate, owing again to the susceptibility of the identification to rewirings. Taken together, these results suggest that caution must be taken when using DWI-based tractography for identifying hubs, as identification is extremely susceptible to false connections.

## Materials and Methods

Data were collected from 10 male adult macaque monkeys (9 *Macaca mulatta*, 1 *Macaca fascicularis*, age 5.8 ± 1.9 years). A subset of these animals (N=3) had MRI-compatible dental acrylic implants anchored to the skull with ceramic bone screws. All surgical and experimental procedures were approved by the Animal Use Subcommittee of the University of Western Ontario Council on Animal Care and were in accordance with the Canadian Council of Animal Care guidelines.

Surgical preparation and anaesthesia protocols have been previously described (Hutchison et al 2011). Briefly, animals were anaesthetized before their scanning session and anaesthesia was maintained using 1.5-2.0% isoflurane during image acquisition. Images were acquired using a 7-T Siemens MAGNETOM head scanner with a high performance gradient (Siemens AC84 II; Gmax = 80 mT/m; SlewRate = 400 T/m/s and an in-house designed and manufactured 8-channel transmit, 24-channel receive coil optimized for the primate head. For each monkey, two diffusion weighted scans were acquired with opposite phase encoding in the superior-inferior direction at 1 mm isotropic resolution. For seven animals data was acquired 2D EPI diffusion (Siemens Advanced Diffusion WIP 511) with TE/TR = 48.8 ms / 7500 ms, b = 1000 s/mm^2^, 64 directions, 104 × 104 matrix, 24 slices, iPat = 3 and bandwidth of 1923 Hz/px. For the remaining 3 animals, a multiband EPI diffusion sequence (Feinberg et al., 2010; Moeller et al., 2010) was used; TE/TR = 47 ms / 6000 ms, multiband =2, b = 1000 s/mm^2^, 64 directions, 128x 128 matrix, 24 slices, iPat = 2, Partial fourier = 5/8 and bandwidth of 1502 Hz/px. A 3D T1w structural reference was collected for all animals using an MP2RAGE (Marques et al., 2010) acquisition at 500 um isotropic resolution; TE/TR = 3.15 ms / 6500 ms, TI1/TI2 = 800 ms / 2700 ms, flip1/flip2 = 4 / 5, 256 × 256 matrix, 128 slices, iPat=2 and 240 Hz/px bandwidth.

Diffusion-weighted image preprocessing was implemented using the FMRIB Software Library toolbox (FSL v5). This consisted of susceptibility-induced distortion correction using FSL’s ‘topup’ (similar to Andersson et al., 2003) and ‘eddy’ (Andersson and Sotiropoulos, 2016) functions, and modeling of multiple fiber directions using FSL’s ‘bedpostx’ function (Behrens et al., 2007). ROI parcellations specified in F99 macaque template space were registered using the Advanced Normalization Tools (ANTS) software package (Avants et al., 2011) to each animal’s T1w anatomical image using a nonlinear registration and then linearly registered to diffusion space. Seed and target ROI masks were defined as the white matter (WM) voxels adjacent to each gray matter (GM) ROI, referred to as the GM-WM interface. An exclusion mask for each seed mask was also created using the GM voxels adjacent to the seed mask. For intrahemispheric tracking, exclusion masks of the opposite hemisphere were also used.

Two distinct parcellation schemes were chosen to match available tract-tracing data. The first (‘Markov-Kennedy’) was an intrahemispheric parcellation of 29 ROIs that matched those contributing to the edge-complete connectivity matrix described in Markov et al (2014) (Fig 1A, top). The second (“CoCo-RM”) was a whole-cortex parcellation of 82 ROIs matching the connectivity matrix described in Shen et al. (2012) (also see Kötter and Wanke, 2005) (Fig 1A, bottom).

Tractography was performed between all ROIs using both parcellation schemes with FSL’s ‘probtrackx2’ function. Parameters used for tracking were: 5000 seeds, 2000 steps, 0.5 mm step length, termination of paths that loop back on themselves and rejection of paths that pass through exclusion mask. Curvature threshold was varied (0.2, 0.4, 0.6, 0.8) and distance correction was toggled on and off. The distance correction option produces a connectivity distribution that is the expected length of the streamlines that cross the voxel multiplied by the number of samples that cross it. In effect, the distance correction option serves to increase the weights of long-distance connections. The “default” parameter combination was considered as that commonly used in human studies (curvature threshold: 0.2; distance correction: off).

A structural connectivity matrix for each parameter set in each parcellation in each animal was generated by taking the number of streamlines detected between each ROI pair and dividing it by the total number of streamlines that were successfully sent from the seed mask (i.e., those that were not rejected or excluded). Each connectivity matrix was subsequently symmetrized.

For distance-based analyses, the distance between ROIs in the Markov-Kennedy parcellation was determined as the geodesic distance of their barycenters, that is, the shortest distance passing through the white matter that connects the barycenters of a pair of areas (as available from the Core-Nets database: core-nets.org). The distance between ROIs in the RM-CoCO parcellation was computed as the Euclidean distance between the centers of all ROIs. For both parcellations, connections were binned according to eight distance quantiles for analysis.

For determining accuracy of the DWI-based structural connectivity matrices (i.e., analyses illustrated in Fig. 2), the corresponding tracer matrices were symmetrized (Fig 1A) and all matrices were binarized. % correct was computed as: # true positives + # true negatives / total # connections (after Azadbakht et al., 2015). We additionally computed the area under the ROC curve (AUC), sensitivity, specificity, and precision of the DWI-derived matrices. Comparisons were only performed on the upper triangles of matrices. Repeated measures one-way ANOVAs were performed to assess the effect of curvature threshold on accuracy, treating the curvature thresholds as four manipulations on the same group of subjects.

To quantify the extent to which physical distance can predict the existence of connections between cortical areas, logistic regression was used, with the existence of connections serving as the binary dependent variable and the physical geodesic distance between the barycenters of cortical areas serving as the independent variable. The model parameters were estimated in a training set consisting of 70% of the total number of area pairs and the rest of the pairs were used to estimate the ROC curve. The procedure was repeated 1000 times and each time the training set was assembled by sampling with replacement. The reported ROC curves and AUC values correspond to these 1000 iterations.

For network analyses, average DWI-based structural connectivity matrices for both the Markov-Kennedy and RM-CoCo parcellations were generated by selecting, for each subject, the matrix having maximum AUC when compared to the tract-tracing data. The combination of tractography parameters and thresholding that yielded the maximum AUC for each subject in each parcellation is provided in Table S2. To enable direct comparison of tracer- and tractography-based networks, tracer matrices were symmetrized. Consequently, network analytic results presented here on the tracer matrices differ slightly from previous studies (e.g., Ercsey-Ravasz et al 2013; Knoblauch et al chapter on RC; Shen et al 2012; Shen et al 2015 RC). Tractography-based networks were then thresholded to match the density of the corresponding tracer-derived networks. In the case of the RM-CoCo parcellation, intra- and interhemispheric quadrants of the tractography-based network were treated with different thresholds due to their vast differences in density in the tract-tracing network (intra: 0.84 vs inter: 0.36). For the Markov-Kennedy parcellation, edges in both tracer- and tractography-based networks were treated as weighted. For the RM-CoCo parcellation, because of the categorical nature of the weighted information in the CoCoMac database, edges were binarized for the tracer-based network as well as the tractography-based network for computing centrality measures (degree and betweenness).

Measures of centrality, modularity partitioning and participation coefficients were obtained using functions from the Brain Connectivity Toolbox (BCT; https://sites.google.com/site/bctnet/). For weighted graphs, the degree of each node was computed as the sum of its edge weights while for binarized graphs, node degree was taken as the total number of its edges.

For the calculation of betweenness centrality, the edge weights were inverted so that larger weights corresponded with longer paths. Community detection was performed using the Louvain algorithm (Blondel et al., 2008). As we did not know how the small network of 29 nodes from the Markov-Kennedy parcellation should be partitioned, we first varied the resolution parameter (gamma) between 0 and 2 in increments of 0.05 and determined the most commonly detected number of partitions >1 in that range. The minimum value of gamma that produced that number of partitions was selected. For the RM-CoCo parcellation, we similarly varied the resolution parameter but selected a gamma value that gave a reasonable number of partitions based on previous studies of whole-brain modularity in the macaque (Harriger et al., 2012; Goulas et al., 2015). Partitioning for both parcellations was then repeated 100 times using the selected gamma value, and the most consistent partitioning was chosen for analysis. This was done independently for both tracer- and tractography-based networks. For the Markov-Kennedy parcellation partitions, both tracer (gamma = 0.65) and tractography (gamma = 0.65) networks were consistently partitioned 100/100 times. For the RM-CoCo parcellation, the most common partition of the tracer network (gamma = 1) occurred 17/100 times while that of the tractography network (gamma = 0.95) occurred 44/100 times. Spearman rank correlation coefficients were computed to assess the correspondence of network measures between the two modalities. Statistical significance of the correlations was assessed using permutation tests by resampling data pairs without replacement 10,000 times.

Rich club detection was performed following the procedures described by Alstott et al. (2014) for computing null networks that are topological, weighted, and of mixed topo-weighted form. Core-periphery detection was performed as described in Ercszey-Ravasz et al (2013). Briefly, the cortico-cortical network was subject to a modified Bron-Kerbosch algorithm with both pivoting and degeneracy ordering (Eppstein et al., 2010). The algorithm detects all cliques up to the maximum size. A clique is a subset of the nodes of the network among which the maximum possible amount of connections exists. The core was defined as the union of all the nodes participating in the cliques of maximum size.

## Acknowledgements

This work was supported by a grant from the Canadian Institutes of Health Research (CIHR) (S.E.) and the J. S. McDonnell Foundation (A.R.M.).

